# A Biodegradable, Tacrolimus-releasing Nerve Wrap Promotes Peripheral Nerve Regeneration

**DOI:** 10.1101/2021.10.23.465561

**Authors:** Simeon C. Daeschler, Katelyn Chan, Konstantin Feinberg, Marina Manoraj, Jenny Cheung, Jennifer Zhang, Kaveh Mirmoeini, J. Paul Santerre, Tessa Gordon, Gregory H. Borschel

## Abstract

Axonal regeneration following nerve repair is slow and often incomplete, resulting in poor functional recovery and sometimes lifelong disability. Yet, there are no FDA-approved therapies available to promote nerve regeneration. Tacrolimus accelerates axonal regeneration, but systemic side-effects presently outweigh its potential benefits for peripheral nerve surgery. We have developed a biodegradable drug delivery system for the sustained local release of tacrolimus at the nerve repair site, with suitable properties for large-scale manufacturing and clinical application, aiming to promote axonal regeneration and functional recovery with minimal systemic drug exposure. Tacrolimus is encapsulated in polycarbonate-urethane nanofibers and electrospun to generate an implantable nerve wrap that releases therapeutic doses of bioactive tacrolimus over 31 days. Size and drug loading are adjustable for applications in small and large caliber nerves, and the wrap degrades within 120 days into biocompatible byproducts. Tacrolimus released from the nerve wrap promotes axon elongation in vitro and accelerates nerve regeneration and functional recovery in preclinical nerve repair models while systemic drug exposure is reduced by 80% compared to systemic delivery. Given its surgical suitability and preclinical efficacy and safety, this system may provide a readily translatable approach to support axonal regeneration and recovery in patients undergoing nerve surgery.

## 1. Introduction

Although the peripheral nervous system is capable of regeneration, functional recovery following nerve repair is slow and often incomplete ^1,2^. Particularly in proximal nerve injuries, the inefficient rate of axonal regeneration of 1 - 2 mm/day ^3,4^ represents a key issue.

Even after meticulous nerve repair the regenerating axons may need to traverse regeneration distances of tens of centimeters (e.g. > 25 cm in adult ulnar nerve injuries at the elbow level for intrinsic hand muscle reinnervation ^5^), resulting in prolonged denervation of the distal nerve and target organ. This leads to Schwann cells atrophy^6^ and occlusion of endoneurial tubes by collagen filaments^7^ rendering the neural pathways increasingly less permissive for reinnervation. Simultaneously, muscle fibers undergo significant cellular changes in response to denervation^8–10^ leading to the progressive disintegration of the contractile apparatus^11,12^ and fibrotic remodeling of the denervated muscle^13,14^. Strategies that can accelerate axonal regeneration may reduce pathway denervation time and increase the number of neurons that reach their targets.

Approximately 43.8 per million people in the United States suffer traumatic nerve injuries annually^16^, most of which affect the upper extremity nerves that control crucial hand functions^16,17^. For the often young patients^18^, these injuries require prolonged rehabilitation^19,20^ with an average time off work of up to 31 weeks^20,21^, affecting mental health^22^ and quality of life^23^. On a societal scale, loss of production and a potential life-long dependency on invalidity benefits cause considerable follow-up costs^1,18,20^ that may exceed $1,000,000 per patient^18^ over their post-injury lifetime.

To date, adjunct pharmaceutical therapies that can promote axonal regeneration following nerve surgery are clinically unavailable. Lacking clinical translation of promising experimental strategies may be attributable to toxicity concerns of bioactive agents necessitating comprehensive safety and efficacy studies. To overcome these concerns, we focused on tacrolimus, an FDA-approved calcineurin inhibitor, which is widely used clinically as an immunosuppressant^24–28^. Independent from its immunosuppressive actions, tacrolimus exhibits direct neurotrophic effects mediated by FKBP52^29–32^ and promotes axonal regeneration in vitro^31,33^ and in vivo by 12% to 16%^34,35^, which translates into faster functional recovery in rodents^34,36,37^. Clinically, the restoration of nerve function beyond anticipated levels was described in patients that received tacrolimus long-term following upper limb transplantation^38,39^ and replantation^40^. However, side effects of systemically delivered tacrolimus, mainly nephrotoxicity^41^, often outweigh the expected benefits for patients undergoing nerve surgery^42^.

Previous studies from our laboratory have shown that the local delivery of tacrolimus to injured axons can promote their regeneration whilst minimizing systemic exposure to other organs^43,44^. We continued this work by developing an implantable local drug delivery system with more suitable properties for clinical translation to enable neuroregenerative therapy at the site of nerve repair. Here we present the system’s physical and drug release properties, demonstrate easy microsurgical applicability, and determine therapeutic efficacy and safety in preclinical models of nerve injury and repair.

## 2. Results

### 2.1. Design and characterization of the local drug delivery system

We chose a nerve wrap design for local tacrolimus delivery. Similar designs are used clinically to prevent scarring around entrapped peripheral nerves by providing a biodegradable interface between nerve and the surrounding tissue^45,46^. Biocompatible polycarbonate urethane (PCNU) synthesized from hexane diisocyanate, poly (hexamethylene carbonate) diol and butanediol served as the encapsulation polymer. PCNU’s degradation products are considered non-toxic ^47^ and PCNU fibers largely retain their mechanical strength and size during biodegradation, which minimizes the risk of nerve compression.

Using coaxial electrospinning, we fabricated a porous PCNU nanofiber matrix to allow for diffusion and cell migration through the construct **(Figure 1A)**. The nanofibers featured a core-shell architecture, with tacrolimus being encapsulated in the inner core to generate a drug reservoir for sustained release **(Figure 1B)**. The electrospun matrix was then sectioned into nerve wraps, irradiation sterilized (25 kGy), and stored at 4 °C until use (Figure 1A). Scanning electron microscopy revealed smooth and non-beaded fibers **(Figures 1B and C)**. Inclusion of tacrolimus did not affect the matrix porosity (*p* = 0.64; **Figure 1E)**, fiber diameter (*p* = 0.62; **Figure 1F)** or strength (*p* = 0.73; **Figure 1G)** when compared to pure polymer fibers.

**Figure 1.**
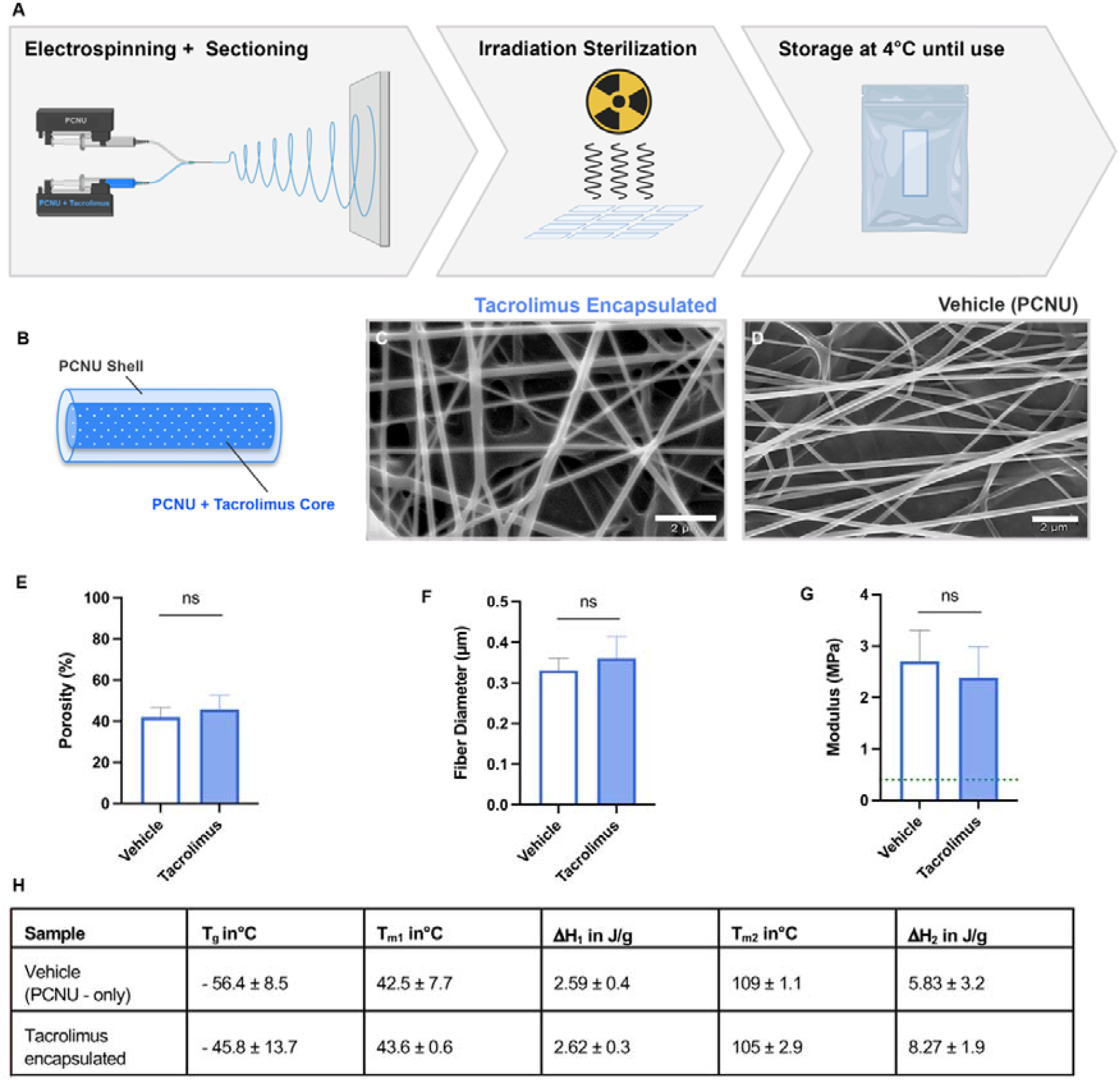
Manufacturing and physical characterization. **(A)** The manufacturing process of the nerve wrap includes co-axial electrospinning, sectioning and gamma irradiation sterilization with subsequent storage at 4°C. **(B)** The nanofiber architecture follows a core-shell principle with tacrolimus being encapsulated in the core polymer to create extended drug release during biodegradation. The uniform fiber morphology of **(C)** - tacrolimus encapsulated- and **(D)** pure polymer nanofibers; no beading was observed (scanning electron microscopy images, 2µm scale bar). **(E)** The matrix porosity and **(F)** average fiber diameter of pure polymer nerve wraps (vehicle, n=8) did not differ from tacrolimus nerve wraps (n=6). **(G)** The dry elastic modulus of the nerve wrap exceeds the tensile strength of human peripheral nerves (dashed line) indicating sufficient strength to withstand surgical manipulation and suturing. **(H)** PCNU has a soft polyol segment and a hard diisocynate/chain extender segment ^48^. Thermal properties of vehicle and tacrolimus nerve wraps including glass transition temperature (T_g_), segment melt temperatures (T_m1_ and T_m2_), and segment melt enthalpies (ΔH_1_ and ΔH_2_), show similar thermal properties with the encapsulation of tacrolimus at the chosen concentration (5.2 w/w%), indicating stability at ambient and body temperature. PCNU – Polycarbonate urethane. Graphs show mean ± SEM.

Clinical application requires the device to withstand surgical manipulation and suturing. We performed tensile testing, showing that the nerve wraps’ strength of 2.70 ± 1.05 MPa exceeded the tensile strength of human peripheral nerves averaging 0.54 MPa ^49^, indicating that the nerve wrap likely withstands manipulation during implantation. Differential scanning calorimetry revealed that tacrolimus encapsulation (5.2 w/w%) did not change the microstructure and thermal properties of the polymer **(Figure 1H)**. Thermogravimetric analysis indicated that the nerve wrap remains in an amorphous state at room temperature (24°C > Tg) and retains stability at physiological temperature (37°C > Tg and 37°C < Tm1, Tm2). This may facilitate scale-up, long-term storage, and clinical application.

### 2.2. The nerve wrap efficiently encapsulates and releases bioactive tacrolimus

Based on our previously published work, we targeted a total drug loading of 200µg tacrolimus per wrap with an extended release of greater than 14 days ^44,50^. We designed two different sizes of the nerve wrap, one representing dimensions for potential clinical application (10 × 15 mm, applicable for a nerve cross sectional area up to 12.5 mm^2^ e.g. forearm ulnar nerve^51^), and one intended for preclinical testing in a rat model (2.5 × 0.5 mm). Mass spectrometry revealed an average drug loading of 223 ± 65 µg, with a mean encapsulation efficiency of 96 ± 8 %, indicating minimal loss of tacrolimus during manufacturing. The 31-days drug release profile in vitro **(Figure 2A)**, yielded a biphasic release profile with a maximum release rate of 11.7 ± 3.8 µg/day within the first 12 days and a subsequent maintenance dose averaging 60.7 ± 37.3 ng/day, sustained over 31 days. As drug encapsulation and release may affect its bioactivity, we conducted a neurite extension assay with release media in cultured embryonic rat dorsal root ganglia (DRGs; **Figure 2B to E**). When compared with control DRGs (media only) the neurons that were incubated in nerve wrap release media from 1- and 31-day timepoints, displayed significantly greater 24 h neurite outgrowth, comparable to a positive control (media containing 50 ng/ml fresh tacrolimus; Figure 2 B). This indicates that the nerve wrap releases bioactive tacrolimus in therapeutic doses for at least 31 days.

**Figure 2.**
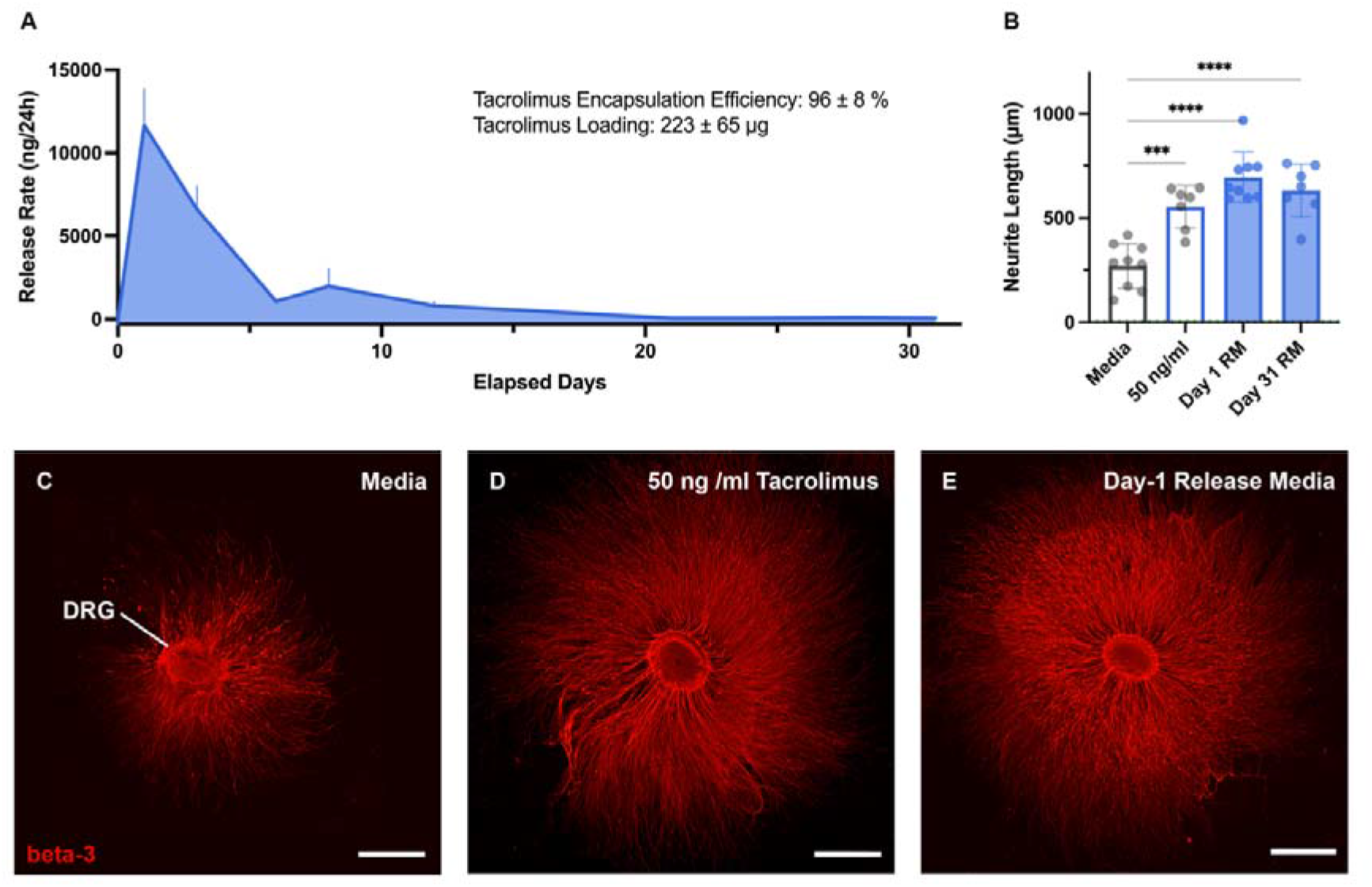
Drug Release and Bioactivity. **(A)** The nerve wraps’ tacrolimus release profile in vitro. **(B)** A neurite extension assay shows significantly longer neurites after 24 h culture when the dorsal root ganglia (DRG) were incubated in day-1 and day-31 nerve wrap release media, compared to a pure media control. No exogenous nerve growth factor (NGF) was added to the cultures. **(C - E)** Representative DRG cultures after 48 h immunostained against the neuronal marker beta 3 tubulin (red) show significantly greater neurite extension in tacrolimus treated cultures (two-tailed t-test: p < 0.05; scale bar 200µm).

### 2.3. The tacrolimus nerve wrap is biocompatible and biodegrades in vivo

Ideally, drug delivery implants degrade after they served their therapeutic purpose, to prevent potential long-term sequelae such as nerve compression. To test for biocompatibility and biodegradability, we implanted the nerve wrap around intact rat sciatic and repaired common peroneal nerves (**Figures 3A and C**). The postoperative swelling of the wrapped nerve segment was significantly reduced when a tacrolimus releasing nerve wrap was implanted as compared to an empty wrap (*p* = 0.03; **Figure 3B**). This may be due to local immunosuppressive effects of tacrolimus potentially improving the device’s biocompatibility. Following implantation, the thickness of the vehicle and tacrolimus nerve wraps decreased to 35.3 ± 10.6 % of the pre-implantation thickness by 7 days, and to 16.7 ± 6.1 % by 60 days **(Figure 3D)**. The wraps were undetectable at 120 days post-implantation, indicating complete biodegradation. Fibrotic capsule formation was not observed.

**Figure 3.**
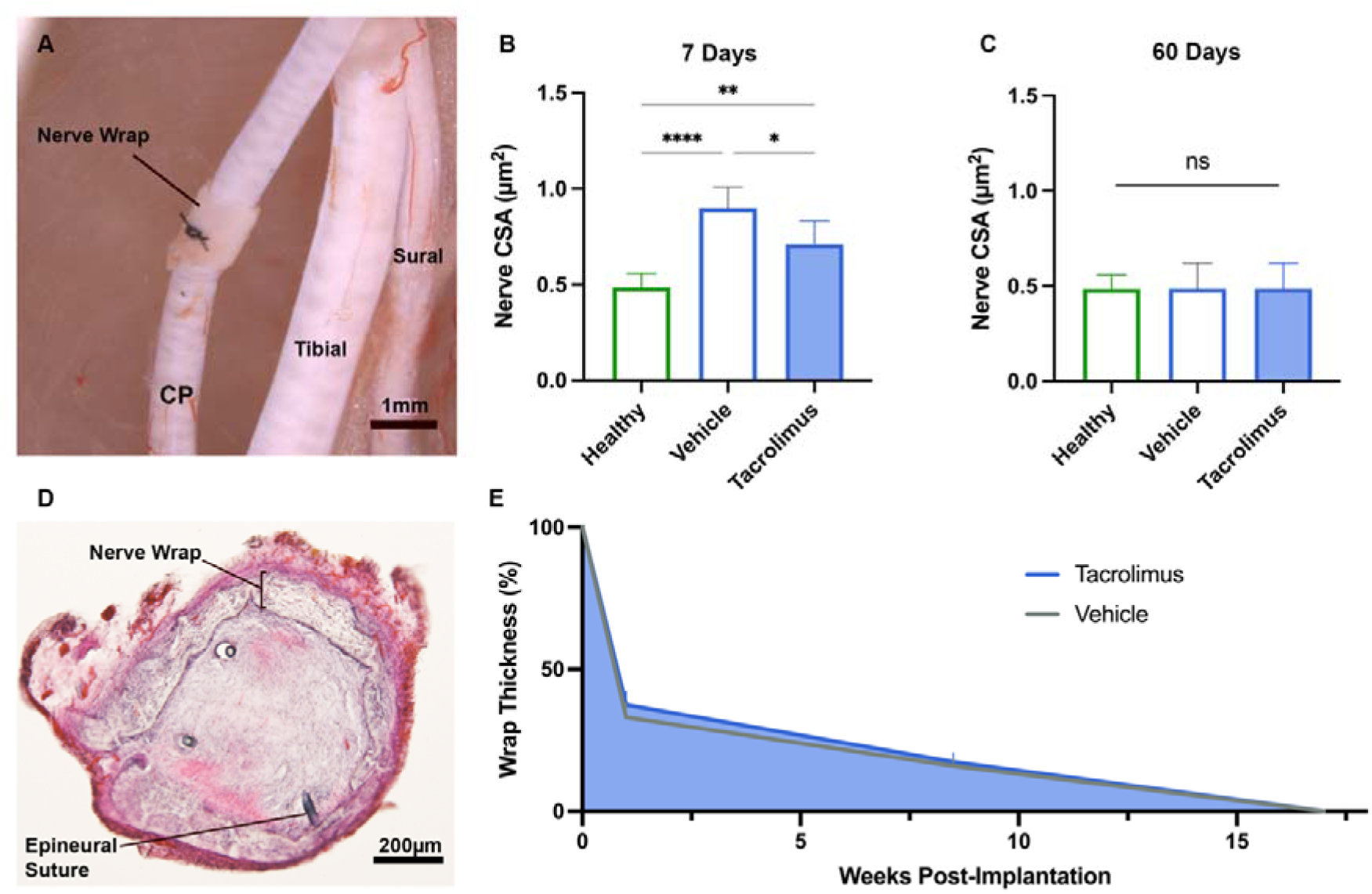
Biocompatibility and -degradation. **(A)** A nerve wrap following implantation around the nerve repair site in a rat the common peroneal nerve. **(B)** Measurements of the nerve cross sectional area following nerve wrap implantation indicate significancy reduced post-operative nerve swelling after tacrolimus nerve wrap implantation as compared to a vehicle nerve wrap. After 60 days post-implantation the nerve cross-sectional area was similar to healthy nerves. **(C)** A cross section through the nerve repair site enclosed by a nerve wrap 7 days post-surgery. **(D)** The biodegradation profile of a tacrolimus nerve wrap (blue) and a vehicle nerve wrap (grey) indicates 65% thinning within the first week following implantation and complete degradation within 120 days. Graphs show mean ± SEM. CP – common peroneal nerve. * p<0.05; **p<0.01; ****p< 0.0001

### 2.4. Tacrolimus nerve wrap implantation increases the number of regenerating nerve fibers following nerve repair

Based on our findings that tacrolimus released form the nerve wrap promoted neurite outgrowth in vitro, we asked whether repaired nerves regenerate more axons when the nerve wrap is implanted around the coaptation site. We transected the common peroneal nerve in rats followed by immediate epineural repair, and either implanted a tacrolimus nerve wrap (200 µg per wrap) or subcutaneously injected tacrolimus daily after surgery (2 mg/kg bodyweight) as previously described^52,53^. Rats that received a vehicle wrap or nerve repair only served as controls. In agreement with the in vitro results (Figure 2B – E), the tacrolimus treated rats regenerated significantly more myelinated nerve fibers 10 mm into the distal nerve segment within three weeks post-surgery when compared to the control groups (**Figure 4E**). Similarly, regenerated axons in nerves that were exposed to the locally delivered tacrolimus were slightly larger as compared to the control groups (**Figure 4F**, p < 0.05), however there was no difference in myelination (**Figure 4G**). Notably, implanting a vehicle nerve wrap had no effect on the number and myelination of nerve fibers compared to nerve repair only (Figure 4E and G, *p* = 0.78 and *p* = 0.98 respectively), indicating that nerve wrap implantation did not cause nerve compression.

**Figure 4.**
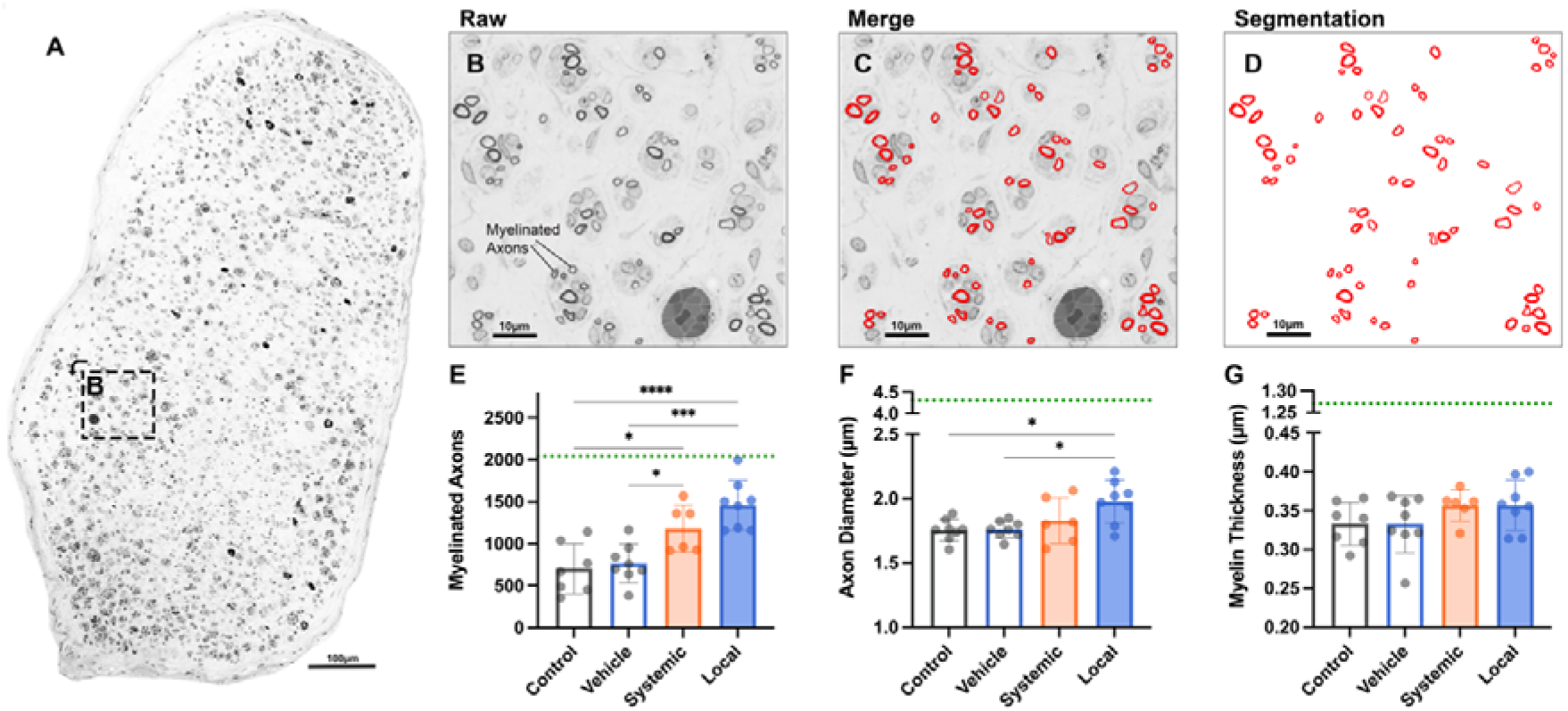
Nerve fiber histomorphometry. **(A**) A representative cross section of a locally tacrolimus treated common peroneal nerve 10 mm distal to the nerve repair site 3 weeks post-surgery. **(B)** In higher magnification multiple thinly myelinated, regenerating nerve fibers are visible. **(C)** A custom-trained deep learning model based on the open-source software AxonDeepSeg^63^ was used to segment entire nerve cross sections in axon / myelin masks (red, **D**) and determine **(E)** the number of myelinated axons, **(F)** axon diameter and **(G)** myelin sheath thickness. Green dotted lines indicate physiological values. * p<0.05; *** p<0.001; **** p<0.0001

### 2.5. Tacrolimus nerve wrap implantation increases the number of regenerating neurons following nerve repair

Next, we asked whether the higher numbers of regenerating nerve fibers in rats that received a tacrolimus nerve wrap reflect in a higher number of regenerating motor and sensory neurons. Three weeks following surgery, we used the fluorescent tracer Fluorogold to retrogradely label neurons that projected their axons at least 7 mm into the distal nerve segment **(Figure 5A to J)**. Consistent with the increased number of regenerated nerve fibers (Figure 4), rats that had received tacrolimus either systemically or locally, had significantly more sensory and motoneurons that regenerated their axons compared to the control groups (**Figure 5K and L**). Application of the vehicle nerve wrap did not affect the number of neurons that regenerated their axons compared to nerve repair only. When analyzing the L3 to L5 DRGs individually, we found that systemically treated rats predominantly regenerated sensory neurons from L3 and L4, whereas in locally treated rats a significantly larger proportion of L5 sensory neurons regenerated their axons (**Figure 5 M**). The cell bodies of L5 neurons being the closest to the drug delivery site, this may suggest the contribution of length-dependent mechanisms (e.g. retrograde axonal transport) in the growth promoting effect of locally delivered tacrolimus.

**Figure 5.**
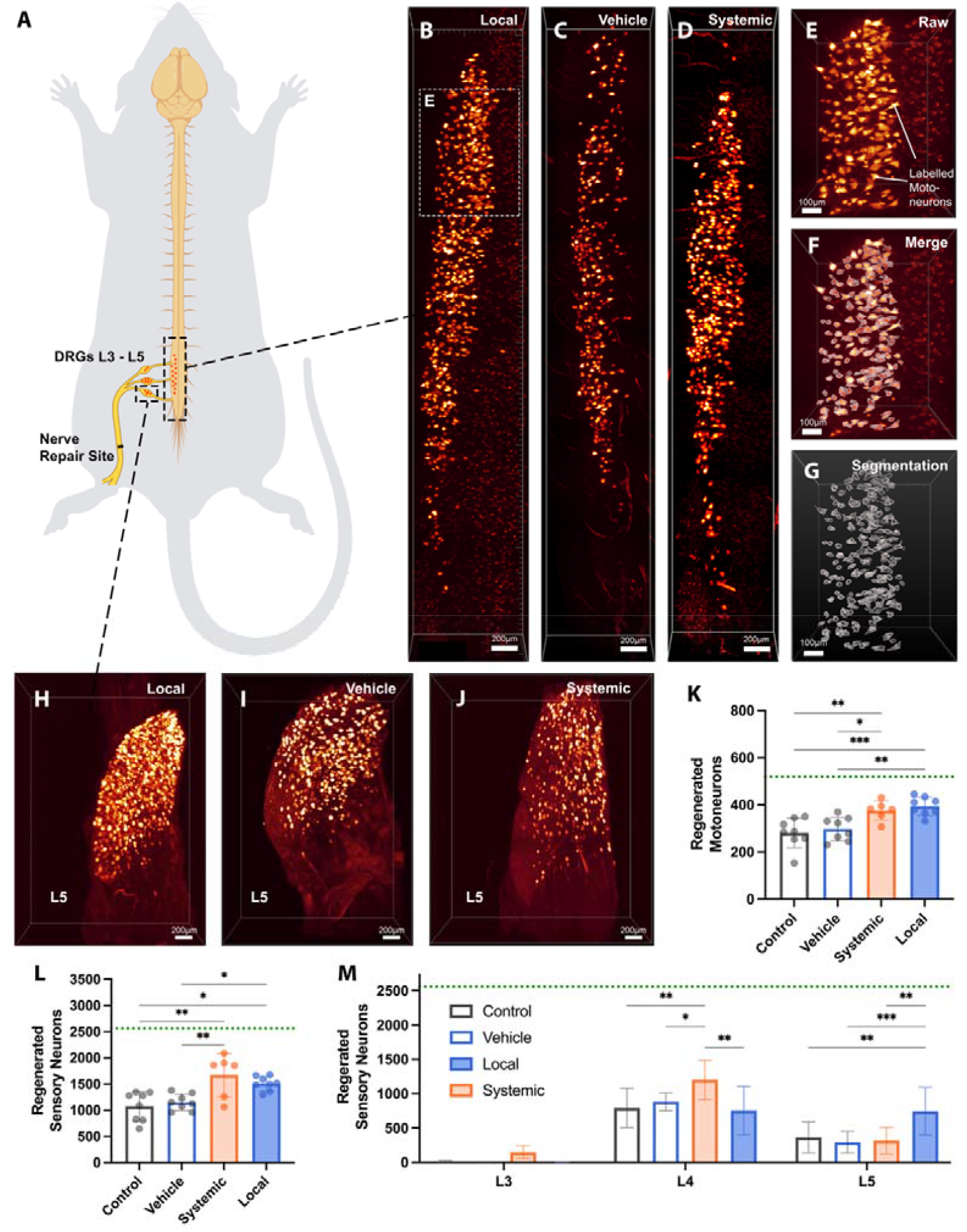
Regenerated motor and sensory neurons. **(A)** Illustration of the localization of back-labelled motor and sensory neurons in relation to the rat common peroneal nerve repair site. **(B)** A representative intact, optically cleared lumbar spinal cord segment of a locally tacrolimus treated rat, **(C)** a vehicle treated rat and **(D)** a systemically treated rat showing fluorogold labelled motoneuron cell bodies (yellow) that project their axons at least 7 mm into the distal nerve segment 3 weeks post-surgery. **(E)** Spinal cord in higher magnification, **(F and G)** with a demonstration of the fluorescence intensity-based neuron segmentation process. **(H)** A representative intact, optically cleared L5 dorsal root ganglion (DRG) of a locally tacrolimus treated rat, **(C)** a vehicle treated rat and **(D)** a systemically treated rat showing fluorogold labelled sensory neuron cell bodies (yellow) that project their axons at least 7 mm into the distal nerve segment 3 weeks post-surgery. **(K)** Quantitation of regenerated motoneurons that project their axons at least 7 mm into the distal nerve segment 3 weeks post-surgery suggests accelerated motor axon regeneration in tacrolimus treated rats. **(L)** Quantitation of regenerated sensory neurons (DRG L3 - L5) that project their axons at least 7 mm into the distal nerve segment 3 weeks post-surgery. Tacrolimus-treated rats regenerated significantly more neurons compared to non-treated and vehicle treated controls. Given the early time point post-surgery, this may indicate accelerated regeneration of sensory axons in vivo. **(M)** DRG level-specific presentation of regenerated sensory neurons showing that rats that received tacrolimus locally at the nerve repair site recruited significantly more neurons from L5 compared to systemically treated and non-treated rats. Green dotted lines indicate the physiological number of neurons that project their axons into intact control common peroneal nerves (mean, n=6). Graphs show means ± SD. Pairwise two-tailed t-tests; * p<0.05; **p<0.01, ***p< 0.001.

### 2.6. Tacrolimus nerve wrap implantation accelerates functional recovery following nerve repair

We then asked whether these changes in neural regeneration result in earlier return of motor function. Clinically, most nerve injures occur in the upper extremity and often affect the median nerve^17^ which is involved in hand motor control. The neuromuscular anatomy of the rat forelimb resembles the human anatomy, with the exception that the rat long finger flexors are exclusively median nerve innervated (**Figure 6A**)^54^. We transected and immediately repaired the median nerve 8 mm proximal to its muscle insertion (flexor digitorum profundus and superficialis) and implanted either a tacrolimus-releasing or a vehicle nerve wrap. Following nerve transection, the rats immediately lost active finger flexion and therefore the ability to grasp. Using daily forepaw grip function tests and video analysis by a blinded investigator (**Figure 6B - E**) we observed that rats that received the tacrolimus nerve wrap required less time to recover active finger flexion when compared to animals that received vehicle wraps (21.4 ± 2.6 days vs 25.2 ± 1.3 days; *p* < 0.05; **Figure 6F and G**). Assuming a 10 mm average regeneration distance from the nerve repair site to the neuromuscular junction, locally delivered tacrolimus accelerated axonal regeneration by 17.6%, or 0.07 mm/day. Differences in grip strength 6 weeks post-surgery did not reach significance (*p* = 0.07; **Figure 6H**).

**Figure 6.**
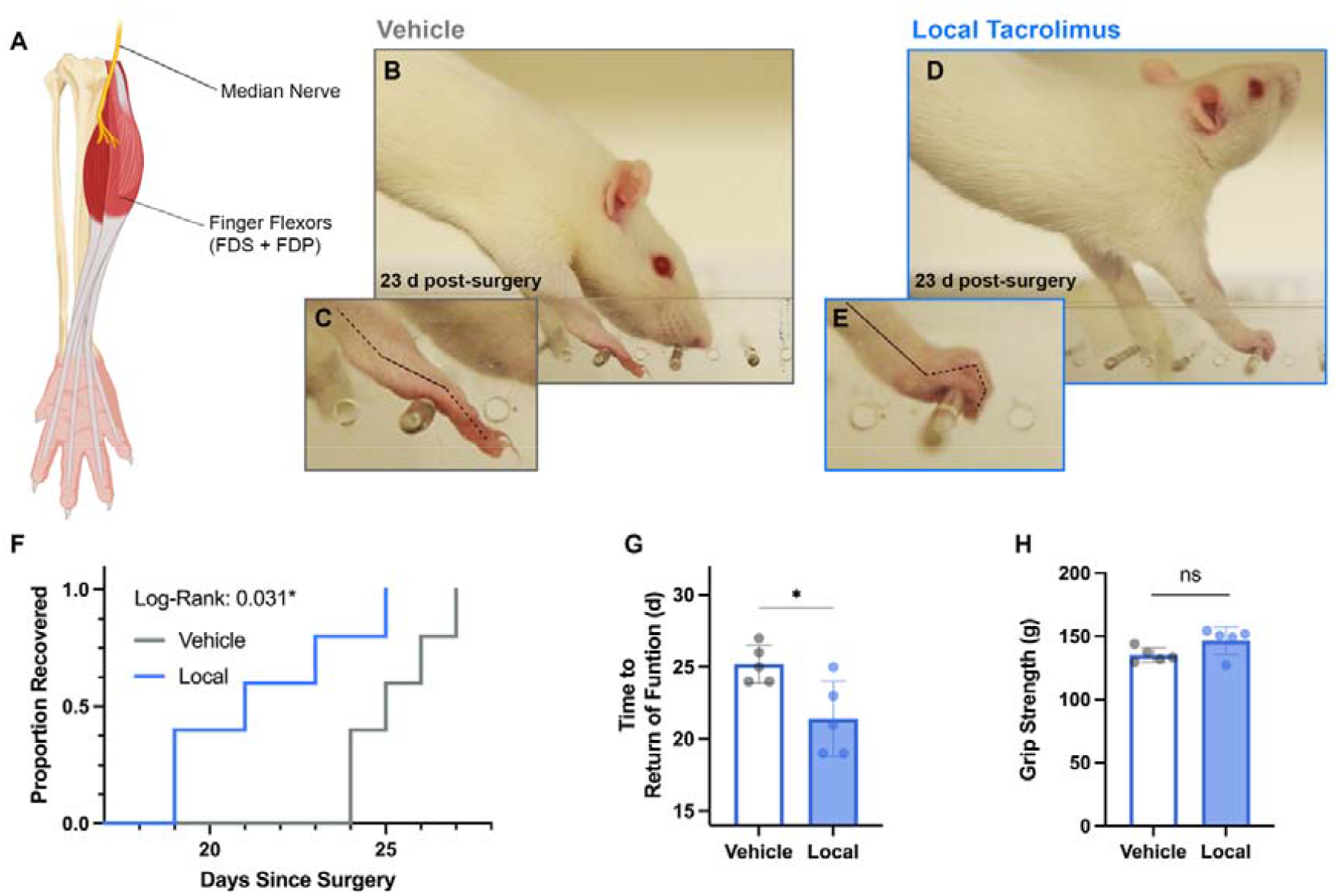
Recovery of Muscle Function. **(A)** Illustration of the rat forelimb anatomy showing the long finger flexors are exclusively median nerve innervated; flexor digitorum profundus (FDP) and - superficialis (FDS). **(B)** Daily grip tests were used to monitor return of active finger flexion following median nerve repair in local tacrolimus and vehicle treated rats. Rats that received a vehicle nerve wrap around the nerve repair site showed no signs of active finger flexion 23-days post-surgery. **(C)** In contrast, 4 of 5 rats that received a tacrolimus releasing nerve wrap around the nerve repair site showed recurring active finger flexion 23-days post-surgery. **(F)** Kaplan Meier curves for motor function recovery of locally tacrolimus treated (n=5) and vehicle treated control rats (n=5) showing significantly accelerated return of motor function in rats that received the tacrolimus nerve wrap following nerve repair. **(G)** Bar graphs showing time to return of active finger flexion in rats that received the tacrolimus or the vehicle nerve wrap following median nerve cut and repair. **(H)** Grip strength of the operated paw 6-weeks post-surgery indicating no significant differences between tacrolimus treated and vehicle treated rats (p = 0.07). * p<0.05; ns = not significant

### 2.7. Implantation of a tacrolimus nerve wrap reduces systemic drug exposure compared to systemic delivery

Systemic exposure to tacrolimus may result in nephro- and liver toxicity^41,55^, and occasionally, neurologic^56^ or psychiatric^57^ symptoms. These side effects presently outweigh the benefits of tacrolimus as a therapy^42^. We used liquid chromatography tandem mass spectrometry to determine the biodistribution and drug exposure of vital organs following local and systemic delivery **(Figure 7A)**. In rats that received the nerve wrap, the tacrolimus concentrations in the kidney, brain, liver, and heart at 7- and 28-days following implantation were significantly lower (−82.2% and −79.0% on average respectively, *p* < 0.05; **Figure 7B and C**) when compared to rats that received tacrolimus systemically (2mg/kg/d). Similarly, the 24h plasma concentration following tacrolimus injection averaged 15.2 ± 11.8 ng/ml with a peak concentration (Cmax) of 30.2 ± 12.4 ng/ml 3h post-injection and a trough concentration (Cmin) of 4.3 ± 0.5 ng/ml 24h post-injection. In contrast, 7 days following tacrolimus nerve wrap implantation, the 24h plasma concentration was significantly lower, averaging 0.3 ± 0.5 ng/ml (p = 0.013) with Cmax of 0.9 ± 0.6 ng/ml and Cmin of 0.09 ± 0.04 ng/ml (**Figure 7 F**). Notably, the tacrolimus concentration in the lumbosacral spinal cord of nerve wrap treated rats, was more than 4-times higher than the average drug concentration in their vital organs 7- and 28-days post-surgery (*p* < 0.05; **Figure 7B to E**) achieving similar drug levels to the systemically treated rats (Figure 7 D and E). Further, the nerve wrap created a significantly higher tacrolimus concentration in the regenerating nerve as compared to the systemic drug delivery at both 7- and 28-days post-surgery (p < 0.01, **Figure 7G and H**). The drug concentration in the contralateral nerve remained low in the locally treated rats whereas the systemically treated rats seemed to accumulate tacrolimus in healthy nerves over time (Figure 7E). These results indicate that the tacrolimus nerve wrap maintains a higher local tacrolimus concentration in the target tissue with reduced systemic exposure compared to systemically delivered tacrolimus.

**Figure 7.**
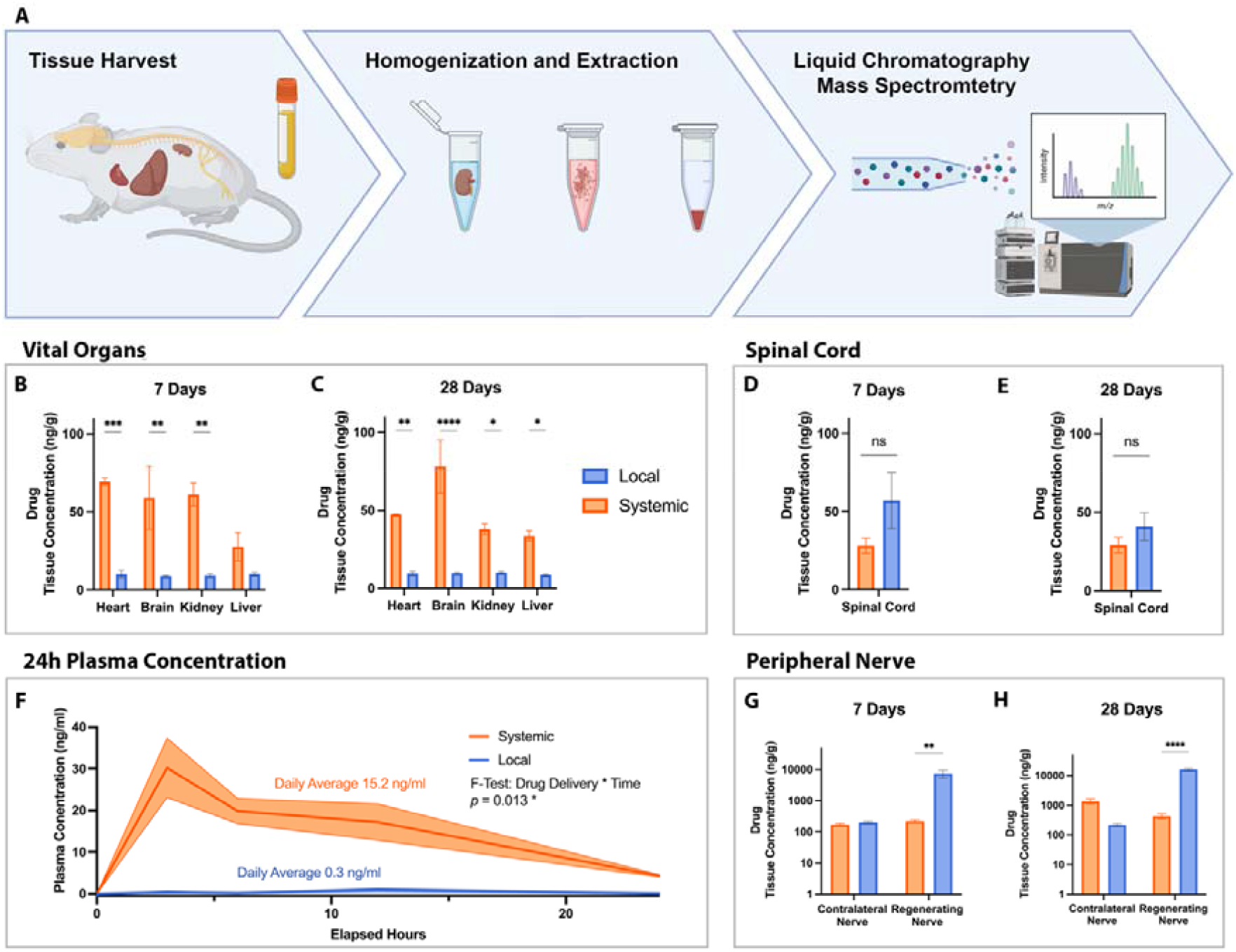
Systemic Drug Exposure Analysis. **(A)** Organs known to be potentially susceptible to tacrolimus exposure, including heart, brain, kidney, and liver, together with neural tissue, including the lumbosacral spinal cord and both common peroneal nerves, and blood plasma were harvested from systemically and locally tacrolimus treated rats 7- and 28-days post-surgery. The snap frozen tissue was homogenized, the tacrolimus was extracted and quantitatively analyzed using liquid chromatography tandem mass spectrometry (LC-MS/MS). **(B)** Tacrolimus tissue concentration in vital organs 7- and **(C)** 28-days post-surgery. Rats that received tacrolimus systemically via daily subcutaneous injections (2mg/kg) showed significantly higher systemic drug exposure compared to rats that received the tacrolimus releasing nerve wrap. **(D)** Tacrolimus tissue concentration in the lumbosacral spinal cord 7- and **(D)** 28-days post-surgery. Local delivery of tacrolimus achieved comparable drug concentrations to systemic tacrolimus delivery (p > 0.05). **(F)** Tacrolimus 24-h plasma concentration profile after subcutaneous injection (2mg/kg) and 7 days post-implantation of the nerve wrap, indicating a significantly reduced circulating mass of tacrolimus when delivered locally. **(G)** Tacrolimus concentration in the regenerating common peroneal nerve and its contralateral counterpart show significantly higher drug concentration in the regenerating nerve 7- and **(H)** 28-days post-surgery when tacrolimus is directly delivered to the nerve repair site. Graphs show mean ± SD; * p<0.05; ** p<0.01; *** p< 0.001; **** p< 0.0001

## 3. Discussion

Here we present a biodegradable drug delivery implant for tacrolimus that enables local and sustained neuroregenerative therapy following peripheral nerve surgery with minimal systemic drug exposure.

We used a polycarbonate urethane (PCNU) polymer platform for its ability to biodegrade in non-cytotoxic byproducts within a clinically reasonable time frame^47,58^. Coaxial electrospinning provided an efficient single-step method to encapsulate tacrolimus within a core-shell nanofiber matrix offering a large surface area-to-volume ratio, with uniform fiber morphology and structural flexibility. A matrix porosity of 40% facilitates diffusion and cell migration^59^ through the construct aiming at minimal interference with the physiological immune response and nerve regeneration process. We validated the design and its drug compatibility by characterizing the morphology and physical properties with and without encapsulated tacrolimus. Scanning electron microscopy showed smooth PCNU-tacrolimus nanofibers and drug encapsulation did not affect fiber morphology. Because of the elastomeric nature of the PCNU, the wrap provided sufficient pliability and tensile strength to minimize pressure along the nerve and yet withstand manipulation during the implantation process. Further, the tacrolimus nerve wrap demonstrated thermal stability at ambient and body temperature thereby facilitating manufacturing, distribution, storage, and clinical use. Based on previously reported work of our laboratory^43,50^ we encapsulated 200µm tacrolimus per wrap into the nanofiber core to generate a sustained drug release^60^. The nanofiber construct achieved higher encapsulation efficiencies of 96% as compared to previously investigated systems^50,61^, indicating minimal drug loss and thus enabling efficient large-scale manufacturing. Further, the construct released therapeutic doses of bioactive tacrolimus for at least 4 weeks, as demonstrated in neurite elongation assays showing over 2.3-fold increased elongation of axons exposed to the nerve wrap release media. Within this time frame, regenerating nerves may be most receptive to local drug delivery at the repair site^62^ as physical barriers including epi- and perineurium as well as the blood-nerve-barrier^63,64^ and myelination^64^ are gradually re-established and thus potentially hinder drug penetration^65^ at later time points. We thus hypothesized a limited therapeutic window for local low-dose tacrolimus therapy early after nerve surgery and produced a biphasic release profile. A release of up to 12 µg/day within the first 12 days immediately yielded a high local drug concentration gradient between the nerve and the surrounding implant, and therefore rapidly attained therapeutic drug levels within the target tissue, followed by a maintenance dose of 60 ng/day on average for up to 31 days. This is corroborated by the drug content analysis via liquid chromatography tandem mass spectrometry showing high tacrolimus concentrations within the treated regenerating nerve 7 days after implantation that are maintained 28 days post-implantation.

For therapeutic efficacy testing of the tacrolimus-releasing nerve wrap we used preclinical models of nerve injury and repair in the rat fore- and hindlimb. Given the relatively short nerve regeneration distances in small animal models we used an early experimental endpoint three weeks post-surgery to capture differences in axonal regeneration rates between treated and control nerves. Rats that received the tacrolimus releasing nerve wrap implanted around the nerve repair site regenerated significantly more axons with a slightly larger diameter compared to control rats. However, higher axon counts may be a result of increased axonal sprouting at the nerve repair site and thus neither necessarily indicate the regeneration of more neurons nor automatically translate into functional benefits. We therefore exposed the regenerated axons to a fluorescent tracer that is retrogradely transported to the neuronal cell body. We found 40% more labelled moto- and 29% more labelled sensory neurons on average in rats that received the tacrolimus releasing nerve wrap following nerve repair as compared to nerve surgery alone. These effect sizes are comparable to rats that were systemically treated with tacrolimus (+ 34% motoneurons and + 43% sensory neurons on average, *p* > 0.05), indicating a similar therapeutic efficacy. Notably, these results represent a snapshot of the early nerve regeneration progress, and thus a higher number of regenerating neurons that regenerate their axons distal to the repair site are most likely the result of accelerated axon elongation as observed in tacrolimus treated neurons in vitro.

To test whether these morphological effects translate into accelerated functional recovery following nerve surgery, we monitored the return of active finger flexion following median nerve transection and repair in rats. We found that rats which received the local drug delivery system regained their ability to grasp 17 % earlier than rats that received an empty drug delivery system. This is in accordance with previously reported effect sizes for systemically delivered tacrolimus on axon elongation in vivo of 12% to 16%^34,35^. Although, this study is limited to one species, and it remains to be determined as to whether this proportion holds true when axons traverse longer regeneration distances and progressively diverge form the drug delivery site, a 17% acceleration may represent weeks of earlier return-of-function for patients undergoing nerve surgery. Given the prolonged disability following upper extremity nerve surgery, therapies that enable earlier return of function may be of significant clinical and socioeconomic benefit.

For application of drug delivery implants in nerve surgery, potential safety concerns include local adverse reactions such as inflammation, fibrotic capsule formation and nerve compression as well as systemic adverse effects including organ toxicity. Drug release systems that completely degrade after serving their therapeutic purpose overcome the need for secondary surgery and reduce the risk of long-term sequalae. Within 7 days following implantation, the wrap thickness decreased by 65% due to hydrolytic degradation, while retaining its macroscopic shape and location for at least 60 days post-implantation before being entirely absorbed within 120 days. However, implant induced local nerve compression may still be a reasonable concern for clinicians, given the susceptibility of axoplasmic transport^66^ and intraneural blood flow^67^ to external pressure.

We therefore compared the neurohistomorphology of repaired nerves after vehicle nerve wrap implantation with repaired nerves without additional treatment. Neither the number of regenerating axons, nor their diameter or myelination state were affected by the empty implant compared to the current gold standard of epineural repair. The local release of tacrolimus further reduced the post-operative nerve swelling by 25% compared to vehicle implantation. This may be attributable to the immunosuppressive effects of tacrolimus potentially supporting the biocompatibility of the implant, counteracting nerve edema and thereby further reducing the risk of local nerve compression. Chronic tissue reactions, such as fibrotic capsule formation, were not observed.

Presently, systemic toxicity is a key limiting factor for tacrolimus in the context of peripheral nerve surgery^42^. By generating a local and sustained, low-dose release profile, the monthly cumulative tacrolimus dose was reduced by more than 98% (200 µg vs 15.5 mg) when compared to previously established systemic drug delivery regimen for promoting nerve regeneration^52,53^. Accordingly, the tacrolimus plasma concentration following local delivery was significantly lower when compared to systemic application, and maximum and average plasma levels remained well below the trough concentration (Cmin) of systemically treated rats. Lower levels of circulating tacrolimus are reflected in the 80% reduced drug exposure of organs that are known to be susceptible to its systemic adverse effects. This indicates that adverse off-target effects may be reduced by local delivery of tacrolimus while its therapeutic efficacy is maintained. For clinical application, one must consider that the volume of distribution in preclinical rodent models is small compared to the human body and thus drug exposure levels are likely to be different. Nevertheless, these results indicate the potential of local tacrolimus delivery to enhance nerve regeneration in clinically relevant effect sizes with minimal systemic drug exposure.

This drug delivery system was designed to provide properties that may facilitate its clinical translation, large-scale manufacturing, and regulatory approval. Future steps towards clinical translation include clinical feasibility studies aiming to capture preliminary information on safety and effectiveness in human subjects. Based on benefit/risk considerations such studies may be initially conducted in patients undergoing primary nerve repair. Evaluation may include longitudinal assessment of tactile thresholds and two-point discrimination, alongside thorough safety testing including pain assessments, regular drug plasma concentration measurements and high-frequency ultrasound imaging to monitor device degradation. Subsequent studies may enroll patients undergoing nerve transfer surgery with its high level of standardization, well-controlled regeneration distances and rapid recovery of motor function allowing clinically meaningful outcome assessments with short trial duration. Two stage procedures such as cross face nerve grafting (stage one) and subsequent free functioning muscle transfer (stage two) after 6 to 12 months may offer the added value of distal nerve graft biopsies at stage two^4^, allowing for detailed histomorphometric analysis of regenerated nerve fibers. Based on these metrics, therapeutic effect sizes can be reliably determined for human subjects and compared to the herein presented preclinical data.

In conclusion, given the extensive clinical safety data for local and systemically applied tacrolimus in various clinical conditions, this bioengineered nanofiber nerve wrap may be a suitable candidate for translation into clinical studies aiming to improve functional outcomes following peripheral nerve surgery.

## 4. Experimental Section/Methods

### Manufacturing and physical characterization of the tacrolimus nerve wrap

Polycarbonate urethane (PCNU) was synthesized of hexane diisocyanate (52649, Sigma Aldrich, Missouri, USA), Poly(hexamethylene carbonate) diol (461172, Sigma Aldrich) and butanediol (309443, Sigma Aldrich) in a 3:2:1 ratio as previously described ^48^. PCNU was mixed with 1,1,1,3,3,3-Hexafluoro-2-propanol (HFIP) (105228, Sigma-Aldrich) to make the polymer solutions for electrospinning, in concentrations of 14 w/v % for the inner core and 20 w/v% for the outer shell. Tacrolimus (LC Laboratories, Massachusetts, USA) was added as a dry powder to inner core polymer solutions at a concentration of 5.2 w/w%. An electrospinning system (NanoSpinner, Inovenso Technology Inc, Massachusetts, USA) was used with a co-axial nozzle of two concentric 22 and 18 gauge blunt-tip needles for the inner core and outer shell, respectively. A flow rate of 0.5 mL/h and a constant voltage difference of 17 kV from nozzle tip to collector (1 kV/cm) was maintained. Electrospun matrices were collected and dried for 72 h under vacuum at room temperature to remove residual solvent. The matrices were then wrapped in sterilization pouches and sterilized by gamma irradiation at a dose of 25 kGy (SOCAAR, University of Toronto, Ontario, CA) and subsequently stored at 4°C until further use. Separate fibrous mats (n=4) were electrospun for each experiment (Figure 2 A - C). Scanning electron microscopy was used for morphological nerve wrap analysis (n=6 tacrolimus encapsulated, n=8 vehicle) from two electrospun matrices. Wraps were mounted, sputter coated with a 15 nm gold layer (Leica EM ACE200, Wetzlar, Germany) and imaged with a FEI XL30 ESEM (Oregon, US) using a beam energy of 20 kV. Average fiber diameter and porosity were determined using the DiameterJ plugin for ImageJ (NIH, https://imagej.nih.gov/ij/). Using pull-to-break tensile testing (840L Servo-All-Electric Test System, TestResources Inc., Minnesota, US) the dry-state elastic modulus was determined for tacrolimus loaded and vehicle matrices (n=3 each) as the slope of the linear region from each plotted stress-strain curve and normalized to the sample thickness.

Thermal gravimetric analyses of tacrolimus and vehicle nerve wraps (n=2 each) were conducted using a thermal gravimetric analyzer (TGA-Q500, TA Instruments, US) under nitrogen gas flux. Samples were heated to 150°C with a ramp rate of 10°C/min and then held under isothermal conditions for 10 min before increasing to a final temperature of 700°C to determine the onset of thermal degradation (T_d_). Differential scanning calorimetry (DSC) was conducted using a TA Instruments Q2000 differential calorimeter (TA Instruments, US) under nitrogen gas flux. Thermograms were recorded between −70 and 200°C at a heating rate of 10°C/min with heat-cool-heat cycles.

### Drug loading, encapsulation efficiency and release profile

Liquid chromatography tandem mass spectrometry (LC-MS/MS) was used to determine tacrolimus loading and encapsulation efficiency for three different tacrolimus loaded matrices. Samples were dissolved in 1 mL HFIP, the solvent was evaporated in a vacuum concentrator (SpeedVac, Thermofisher, MA, US) and the remaining drug was reconstituted in 1 mL acetonitrile (1401-7-40, Caledon Laboratories, Ontario, Canada) with Cyclosporine A as an internal standard, and analyzed as previously described ^43,50^. Briefly, a mass spectrometer (API4000, SCIEX, Ontario, Canada) with a spherisorb column (30×4.6mm; 5 μm; Massachusetts, USA) was used with a mobile phase (65% aqueous acetonitrile, 2 mM ammonium acetate and 0.1% (v/v) formic acid). Quantifications were based on ammonium□adduct transition masses and drug loading was determined as tacrolimus mass per nerve wrap mass. Encapsulation efficiency was defined as ratio of detected tacrolimus loading over the theoretical maximum loading. Drug release was determined from six nerve wraps (10×15mm) derived from three different matrices. Samples were incubated in sterile phosphate buffered saline (PBS) in a water bath shaker (37°C, 50 RPM) and the solution was collected for tacrolimus content analysis via LC-MS/MS. Drug release was calculated as released tacrolimus mass per sample per day.

### Bioactivity testing

Rat dorsal root ganglia (DRG) neurite extension assays were used. Tacrolimus nerve wraps (n=8; 10×15mm) from four different matrices were incubated in Neurobasal media (NBM, 21103049, Thermo Fisher, Massachusetts, USA), 2 vol.% B-27 serum free supplement, 1 vol% penicillin-streptomycin, and 1 vol% L-glutamine. A total of n=16 DRGs were harvested from E15 rat embryos, placed in a Matrigel coated well (Corning, New York, USA) and subjected to four different experimental conditions: Day-one nerve wrap release media (n=4), day-31 release media (n=4), a negative control (NBM only, n=4) and a positive control (50 ng /ml tacrolimus in NBM, n=4). Following 24 hours of incubation, cultures were imaged using phase contrast microscopy (AxioVert, Carl Zeiss Microscopy GmbH, Jena, Germany). Mean neurite extension was measured using ImageJ ^50^. After 48h cultures were fixed in 4% paraformaldehyde (PFA), immunostained against a neuronal marker (rabbit anti-beta-3-tubulin; 18207, Abcam, Cambridge, UK; 1:500 dilution; goat anti-rabbit Alexa Fluor 555 conjugated secondary antibody; A-21428, Invitrogen, California, USA; 1:1000) and imaged using confocal fluorescence microscopy (DMI8, Leica, Wetzlar, Germany).

### In vivo Efficacy and safety testing in an experimental rat model

#### Study design

All animal experiments are reported in accordance with the ARRIVE guidelines ^68^. The primary objectives of the animal experiments were to determine therapeutic efficacy and drug biodistribution of locally and systemically delivered tacrolimus following nerve surgery. The number of rats used in each experimental group was determined by power analysis based on previous experience with rodent nerve injury models and local drug delivery ^43^. For labelled neuron quantifications we considered d=1.5 for motoneurons (assuming a typical SD of 50) and d=2.0 for sensory neurons (assuming an SD of 250) to be minimal relevant effect sizes. To achieve a power of 0.8 on an alpha level of 0.05 (normally distributed, two tailed t-test) this required n=6 and n=9 animals per group respectively. For functional tests we considered d=2.5 to be worth being reported, corresponding to n=4 animals per group. Animals were randomly allocated to experimental groups and the investigators were blinded for outcome assessments. No experimental animals or outliers were excluded from the analysis.

#### Experimental animals

A total of n=76 adult (250 - 300g), female rats with a genetic Sprague Dawley background were included. All animals were housed in a central animal care facility with fresh water and pellet food ad libitum. A constant room temperature (22°C) and a circadian rhythm of 12h per 24h illumination were automatically maintained. All procedures were performed in strict accordance with the National Institutes of Health guidelines, the Canadian Council on Animal Care (CCAC) and were approved by the Hospital for Sick Children’s Laboratory Animal Services Committee.

#### Hindlimb nerve injury and repair model

A total of n=30 rats underwent unilateral common peroneal nerve cut and immediate epineural repair surgery. All surgical procedures were performed under aseptic conditions and inhalation anesthesia with an Isoflurane (Baxter, Illinois, USA) oxygen mixture and analgesia (4 mg/kg body weight extended-release Metacam, Boehringer Ingelheim, Ingelheim, Germany). The common peroneal nerve was exposed through a dorsolateral–gluteal muscle-splitting incision, cut 3 mm distal to the sciatic nerve bifurcation with micro scissors, readapted and repaired using 3 epineural single stitch sutures (10-0 Ethilon, Ethicon, Ohio, USA). After nerve repair, the rats were allocated to the experimental groups (local tacrolimus n=8, local vehicle n=8, systemic tacrolimus n=6, repair-only control n=8) using permuted block randomization. In the local tacrolimus group, a 2.5 × 0.5 mm nerve wrap loaded with 200µg tacrolimus was loosely wrapped around the nerve repair site and held in place by a single stitch (10-0 Ethilon, Ethicon, Ohio, US). The vehicle group received an empty drug delivery system (PNCU polymer only) in an otherwise identical fashion. A repair-only control group received no additional treatment after nerve repair, and systemic tacrolimus group (positive control) received daily subcutaneous injections of tacrolimus (2 mg / kg body weight). The wound was closed in three layers with 5-0 Vicryl (Ethicon, Ohio, US) sutures, and all operated animals were allowed to recover in a warm environment prior to returning to the housing facility.

#### Preparation and systemic delivery of tacrolimus

For systemic delivery, a 10 mg/ml stock solution was prepared by dissolving tacrolimus (LC Laboratories, Woburn, US) in 80% ethanol / 20% Kolliphor (C5135, Sigma Aldrich, Missouri, US), stored at −20°C until use and replaced every 3 days ^53^. Immediately before injection, the stock solution was diluted with sterile double distilled water to a final concentration of 2 mg/ml. Post-surgery, the rats received 2mg/kg tacrolimus daily via subcutaneous injection ^52,53^.

#### Nerve fiber histomorphometry

Three weeks after nerve repair, a common peroneal segment 10 mm distal to the nerve repair site was harvested, immersed in 2.5% glutaraldehyde fixative (G6257, Sigma Aldrich) / 0.1 M sodium cacodylate trihydrate (C0250, Sigma Aldrich) overnight at 4°C and postfixed in 2% osmium tetroxide (75632, Sigma Aldrich) for 2 h. Samples were dehydrated in ascending ethanol series, embedded in epoxy (45345, Sigma Aldrich), sectioned into 1-μm cross-sections (ultramicrotome EM UC7, Leica Microsystems) and imaged (Axiovert 200M, Carl Zeiss Microscopy GmbH, Jena, Germany) using a 63x/1.4 oil objective. A custom-trained deep learning model based on the open-source software *AxonDeepSeg* ^*69*^ determined the number of myelinated nerve fibers, axon diameter and myelin sheath thickness for entire cross sections.

#### Quantitation of regenerated motor and sensory neurons

Three weeks after nerve repair, the common peroneal nerve was cut 7 mm distal to the nerve repair site and the end of the proximal common peroneal nerve segment was placed in a well containing 5 µl 4% Fluoro-Gold (Fluorochrome LLC, Denver, US) in double distilled water. The wound bed was draped to prevent leakage of the florescent tracer. After 60 min of exposure, the wounds were thoroughly irrigated with saline, dried and closed in three layers with 5-0 Vicryl (Ethicon, Ohio, US) sutures. Seven days after the procedure, the spinal cord along with the ipsilateral DRG level L3 to L5 were harvested ^70^, fixed by immersion in 4% PFA at 4°C for 24h, and optically cleared using a modified FDISCO protocol ^71,72^. Briefly, the tissue was immersed in an ascending series of precooled Tetrahydrofuran (186562, Sigma-Aldrich) in double distilled water solutions (50 vol%, 75 vol%, 3 × 100 vol%) adjusted to pH 9.0 with Triethylamine (471283, Sigma-Aldrich). After a subsequent step in 100 % Dichloromethane (270997, Sigma-Aldrich) the specimen was immersed in Ethanol / Dibenzyl ether (108014, Sigma-Aldrich) solutions for refractive index matching ^71,73^. For image acquisition, we used a 3D light-sheet fluorescence microscope (Zeiss Lightsheet Z.1, Carl Zeiss Microscopy GmbH, Jena, Germany) equipped with a 405 nm (20mW) laser, a 20x / 1.0 CLARITY objective and a p.co edge 5.5 Camera (PCO AG, Kehlheim, Germany). For data processing, segmentation, and quantitative analyses, we used an Arivis vision 4D (version 3.0, arivis AG, Rostock, Germany) and Imaris (Version 9.5.1, Bitplane AG, Zurich, Switzerland). Selected images in this publication were created with BioRender.com.

#### Recovery of motor function

Following unilateral median nerve cut and immediate epineural repair, the time to onset of active finger flexion and grip strength at six weeks post-surgery were determined in 10 rats. Perioperative management was as described for the hindlimb model. The right median nerve was exposed by a medial incision along the medial bicipital groove, traced distally to its insertion into the forearm flexors and transected 8 mm proximal to its muscle insertion point, followed by immediate epineural repair. Rats were block randomized in two groups, receiving either a tacrolimus releasing nerve wrap, or an empty nerve wrap (vehicle) as described above. Starting 7 days after surgery, the rats were tested daily for recovery of active long finger flexion using a horizontal ladder grasping test (Figure 6 B - E). The test was repeated 10 times per animal and session to reduce the risk of false negatives, and video-taped for post hoc analysis by a blinded investigator. Active flexion of the proximal interphalangeal joint was considered a positive response. Six weeks after nerve repair, the animals performed a grip strength test as previously described ^74^. Briefly, the rat was held at the base of its tail and a grip response was induced with the post-operative paw on a scale-coupled grip device. Then the rat was pulled away from the grid in a steady, upright motion and the maximum grip strength was determined as the average of the top 3 out of 10 repetitions to account for rounds with submaximal voluntary grip response.

#### Systemic drug exposure analysis

Systemic drug exposure was determined via LC-MS/MS. Twelve rats underwent unilateral common peroneal nerve repair and received either the tacrolimus releasing nerve wrap (n=6) or daily tacrolimus injections (n=6) as described. Seven- and 28-days post-surgery the kidney, brain, liver, heart, the lumbar segment of the spinal cord and both common peroneal nerves were harvested from 3 animals per group and snap frozen in liquid nitrogen. Tissues were weighed, immersed in 300 µl lysis buffer containing 50 mM Tris (17926, Thermo Fisher Scientific, Massachusetts, US), 150 mM NaCl, 2 mM EDTA (E9884, Sigma Aldrich), 0.1% Sodium Dodecyl Sulfate (L3771, Sigma Aldrich), 1% NP40 (Sigma Aldrich), 0.5% DOC and 1% Protease inhibitor and homogenized by sonication on ice for 30 seconds twice ^43^. Tacrolimus was extracted with acetonitrile (1401-7-40, Caledon Laboratories, Ontario, Canada). The solution was homogenized for 30 seconds on ice twice, centrifuged at 4 °C for 5 min at 16,000 G and the supernatant was extracted for analysis with LC-MS/MS ^43^.

#### Biodegradation and biocompatibility in vivo

A total of n=32 sterile nerve wraps (n=16 tacrolimus loaded and n=16 vehicle-only) were implanted around the uninjured sciatic nerve or repaired common peroneal nerve in n=24 randomly allocated rats. The rats were further block randomized into three sub-groups of n=4 rats each for analysis 7-, 60- or 120-days post-implantation. The wrapped nerve segment (8-10 mm) was harvested, immersion fixed in 4% precooled PFA overnight at 4°C, cryoprotected in 30% sucrose / 4% PFA solution for 3 to 5 days, embedded in tissue freezing medium (23-730-571, Fisher Scientific, Massachusetts, USA), snap frozen with liquid nitrogen and stored in −80°C. Nerve cross sections (20 µm) were cut over the entire length (CM3050S cryostat, Leica Biosystems, Nuβloch, Germany), slide mounted and Hematoxylin & Eosin stained. Sections were imaged at 20X magnification (Panoramic 250 Flash II Slide Scanner, 3DHistech, Budapest, Hungary) and CaseViewer software (3DHistech, Budapest, Hungary) was used for measurements of nerve cross sectional area and wrap thickness.

### Statistical analysis

We used JMP (version 15.1.0, SAS Institute, Cary, USA) and GraphPad Prism (GraphPad Software, San Diego, California USA) for statistical analysis. Descriptive statistics were calculated, and means are expressed with standard deviations (± SD). To test for normality of continuous variables, we used normal quantile plots and Anderson-Darling tests. For physical nerve wrap characterization, one-way analysis of variance (ANOVA) was conducted. For *in vivo* safety and efficacy testing we used two-sided Student’s t tests for pairwise comparison of normally distributed, continuous data. For non-normally distributed variables we used Wilcoxon tests. For repeated measure comparison such as drug plasma concentration 24 h profile, we used a multivariate repeated-measures approach with multivariate F-tests. A significance level of 5% was used (*p* < 0.05).

## Acknowledgments

The authors would like to thank Matthew W. Forbes, PhD, Department of Chemistry, University of Toronto for his advice and support with the mass spectrometry experiments. The authors would like to acknowledge the members of the J. Paul Santerre Lab at the University of Toronto, who helped with the synthesis of polycarbonate urethane and shared their expertise. The authors would like to thank Meghan McFadden, PhD, Ted Rogers Centre for Heart Research, for assistance with mass spectrometry sample preparation.

This work was supported by the German Research Foundation (DA 2255/1-1), a SickKids Research Training Competition (RESTRACOMP) Graduate Scholarship, an Ontario Graduate Scholarship, Natural Sciences and Engineering Research Council of Canada (NSERC) and a Kickstarter grant from the Institute of Biomedical Engineering (BME) at the University of Toronto. SCD and KC contributed equally to this work

## References

1 Jaquet, J. B. et al. Median, ulnar, and combined median-ulnar nerve injuries: functional outcome and return to productivity. J Trauma 51, 687–692 (2001).

2 Lan, C. Y. et al. Prognosis of Traumatic Ulnar Nerve Injuries: A Systematic Review. Annals of plastic surgery 82, S45–s52, doi:10.1097/sap.0000000000001727 (2019).

3 Seddon, H. J., Medawar, P. B. & Smith, H. Rate of regeneration of peripheral nerves in man. The Journal of physiology 102, 191–215 (1943).

4 Braam, M. J. & Nicolai, J. P. Axonal regeneration rate through cross-face nerve grafts. Microsurgery 14, 589–591, doi:10.1002/micr.1920140909 (1993).

5 Paulos, R. & Leclercq, C. Motor branches of the ulnar nerve to the forearm: an anatomical study and guidelines for selective neurectomy. Surg Radiol Anat 37, 1043–1048, doi:10.1007/s00276-015-1448-1 (2015).

6 Terenghi, G., Calder, J. S., Birch, R. & Hall, S. M. A morphological study of Schwann cells and axonal regeneration in chronically transected human peripheral nerves. Journal of hand surgery (Edinburgh, Scotland) 23, 583–587, doi:10.1016/s0266-7681(98)80006-5 (1998).

7 Röyttä, M. & Salonen, V. Long-term endoneurial changes after nerve transection. Acta neuropathologica 76, 35–45, doi:10.1007/bf00687678 (1988).

8 Goldspink, D. F. The effects of denervation on protein turnover of the soleus and extensor digitorum longus muscles of adult mice. Comparative Biochemistry and Physiology Part B: Comparative Biochemistry 61, 37–41, doi:https://doi.org/10.1016/0305-0491(78)90210-9 (1978).

9 Cisterna, B. A. et al. Active acetylcholine receptors prevent the atrophy of skeletal muscles and favor reinnervation. Nature communications 11, 1073, doi:10.1038/s41467-019-14063-8 (2020).

10 Cea, L. A. et al. De novo expression of connexin hemichannels in denervated fast skeletal muscles leads to atrophy. Proceedings of the National Academy of Sciences of the United States of America 110, 16229–16234, doi:10.1073/pnas.1312331110 (2013).

11 Lu, D. X., Huang, S. K. & Carlson, B. M. Electron microscopic study of long-term denervated rat skeletal muscle. The Anatomical record 248, 355–365, doi:10.1002/(sici)1097-0185(199707)248:3<355::Aid-ar8>3.0.Co;2-o (1997).

12 Boncompagni, S. et al. Structural differentiation of skeletal muscle fibers in the absence of innervation in humans. Proceedings of the National Academy of Sciences of the United States of America 104, 19339–19344, doi:10.1073/pnas.0709061104 (2007).

13 Borisov, A. B., Huang, S. K. & Carlson, B. M. Remodeling of the vascular bed and progressive loss of capillaries in denervated skeletal muscle. The Anatomical record 258, 292–304, doi:10.1002/(sici)1097-0185(20000301)258:3<292::Aid-ar9>3.0.Co;2-n (2000).

14 Gauthier, G. F. & Dunn, R. A. Ultrastructural and cytochemical features of mammalian skeletal muscle fibres following denervation. Journal of cell science 12, 525–547 (1973).

15 Ruijs, A. C., Jaquet, J. B., Kalmijn, S., Giele, H. & Hovius, S. E. Median and ulnar nerve injuries: a meta-analysis of predictors of motor and sensory recovery after modern microsurgical nerve repair. Plastic and reconstructive surgery 116, 484–494; discussion 495-486, doi:10.1097/01.prs.0000172896.86594.07 (2005).

16 Karsy, M. et al. Trends and Cost Analysis of Upper Extremity Nerve Injury Using the National (Nationwide) Inpatient Sample. World Neurosurg 123, e488–e500, doi:10.1016/j.wneu.2018.11.192 (2019).

17 Tapp, M., Wenzinger, E., Tarabishy, S., Ricci, J. & Herrera, F. A. The Epidemiology of Upper Extremity Nerve Injuries and Associated Cost in the US Emergency Departments. Annals of plastic surgery 83, 676–680, doi:10.1097/sap.0000000000002083 (2019).

18 Hong, T. S. et al. Indirect Cost of Traumatic Brachial Plexus Injuries in the United States. J Bone Joint Surg Am 101, e80, doi:10.2106/jbjs.18.00658 (2019).

19 Rosberg, H. E. et al. Injury to the human median and ulnar nerves in the forearm--analysis of costs for treatment and rehabilitation of 69 patients in southern Sweden. Journal of hand surgery (Edinburgh, Scotland) 30, 35–39, doi:10.1016/j.jhsb.2004.09.003 (2005).

20 Bergmeister, K. D. et al. Acute and long-term costs of 268 peripheral nerve injuries in the upper extremity. PloS one 15, e0229530, doi:10.1371/journal.pone.0229530 (2020).

21 Bruyns, C. N. et al. Predictors for return to work in patients with median and ulnar nerve injuries. The Journal of hand surgery 28, 28–34, doi:10.1053/jhsu.2003.50026 (2003).

22 Yannascoli, S. M., Stwalley, D., Saeed, M. J., Olsen, M. A. & Dy, C. J. A Population-Based Assessment of Depression and Anxiety in Patients With Brachial Plexus Injuries. The Journal of hand surgery 43, 1136.e1131-1136.e1139, doi:10.1016/j.jhsa.2018.03.056 (2018).

23 Ciaramitaro, P. et al. Traumatic peripheral nerve injuries: epidemiological findings, neuropathic pain and quality of life in 158 patients. Journal of the peripheral nervous system : JPNS 15, 120–127, doi:10.1111/j.1529-8027.2010.00260.x (2010).

24 Haddad, E. M. et al. Cyclosporin versus tacrolimus for liver transplanted patients. The Cochrane database of systematic reviews, Cd005161, doi:10.1002/14651858.CD005161.pub2 (2006).

25 Webster, A., Woodroffe, R. C., Taylor, R. S., Chapman, J. R. & Craig, J. C. Tacrolimus versus cyclosporin as primary immunosuppression for kidney transplant recipients. The Cochrane database of systematic reviews, Cd003961, doi:10.1002/14651858.CD003961.pub2 (2005).

26 Cury Martins, J. et al. Topical tacrolimus for atopic dermatitis. The Cochrane database of systematic reviews 2015, Cd009864, doi:10.1002/14651858.CD009864.pub2 (2015).

27 Lee, Y. H. et al. Tacrolimus for the treatment of active rheumatoid arthritis: a systematic review and meta-analysis of randomized controlled trials. Scand J Rheumatol 39, 271–278, doi:10.3109/03009740903501642 (2010).

28 Baumgart, D. C., Macdonald, J. K. & Feagan, B. Tacrolimus (FK506) for induction of remission in refractory ulcerative colitis. The Cochrane database of systematic reviews, Cd007216, doi:10.1002/14651858.Cd007216 (2008).

29 Gold, B. G., Densmore, V., Shou, W., Matzuk, M. M. & Gordon, H. S. Immunophilin FK506-binding protein 52 (not FK506-binding protein 12) mediates the neurotrophic action of FK506. The Journal of pharmacology and experimental therapeutics 289, 1202–1210 (1999).

30 Daneri-Becerra, C., Patiño-Gaillez, M. G. & Galigniana, M. D. Proof that the high molecular weight immunophilin FKBP52 mediates the in vivo neuroregenerative effect of the macrolide FK506. Biochem Pharmacol 182, 114204, doi:10.1016/j.bcp.2020.114204 (2020).

31 Quintá, H. R. & Galigniana, M. D. The neuroregenerative mechanism mediated by the Hsp90-binding immunophilin FKBP52 resembles the early steps of neuronal differentiation. Br J Pharmacol 166, 637–649, doi:10.1111/j.1476-5381.2011.01783.x (2012).

32 Steiner, J. P. et al. Neurotrophic actions of nonimmunosuppressive analogues of immunosuppressive drugs FK506, rapamycin and cyclosporin A. Nature medicine 3, 421–428, doi:10.1038/nm0497-421 (1997).

33 Lyons, W. E., George, E. B., Dawson, T. M., Steiner, J. P. & Snyder, S. H. Immunosuppressant FK506 promotes neurite outgrowth in cultures of PC12 cells and sensory ganglia. Proceedings of the National Academy of Sciences 91, 3191, doi:10.1073/pnas.91.8.3191 (1994).

34 Udina, E., Ceballos, D., Verdú, E., Gold, B. G. & Navarro, X. Bimodal dose-dependence of FK506 on the rate of axonal regeneration in mouse peripheral nerve. Muscle & nerve 26, 348–355, doi:10.1002/mus.10195 (2002).

35 Gold, B. G., Katoh, K. & Storm-Dickerson, T. The immunosuppressant FK506 increases the rate of axonal regeneration in rat sciatic nerve. The Journal of neuroscience : the official journal of the Society for Neuroscience 15, 7509–7516, doi:10.1523/jneurosci.15-11-07509.1995 (1995).

36 Gold, B. G., Storm-Dickerson, T. & Austin, D. R. The immunosuppressant FK506 increases functional recovery and nerve regeneration following peripheral nerve injury. Restorative neurology and neuroscience 6, 287–296, doi:10.3233/rnn-1994-6404 (1994).

37 Sulaiman, O. A., Voda, J., Gold, B. G. & Gordon, T. FK506 increases peripheral nerve regeneration after chronic axotomy but not after chronic schwann cell denervation. Experimental neurology 175, 127–137, doi:10.1006/exnr.2002.7878 (2002).

38 Schuind, F. et al. The first Belgian hand transplantation--37 month term results. Journal of hand surgery (Edinburgh, Scotland) 31, 371–376, doi:10.1016/j.jhsb.2006.01.003 (2006).

39 Owen, E. R. et al. Peripheral nerve regeneration in human hand transplantation. Transplant Proc 33, 1720–1721, doi:10.1016/s0041-1345(00)02656-7 (2001).

40 Martin, D. et al. [First case in the world of autoreplantation of a limb associated with oral administration of an immunosupressant agent (FK 506-Tacrolimus)]. Ann Chir Plast Esthet 50, 257–263, doi:10.1016/j.anplas.2005.02.001 (2005).

41 Bentata, Y. Tacrolimus: 20 years of use in adult kidney transplantation. What we should know about its nephrotoxicity. Artif Organs 44, 140–152, doi:10.1111/aor.13551 (2020).

42 Zuo, K. J., Saffari, T. M., Chan, K., Shin, A. Y. & Borschel, G. H. Systemic and Local FK506 (Tacrolimus) and its Application in Peripheral Nerve Surgery. The Journal of hand surgery 45, 759–765, doi:10.1016/j.jhsa.2020.03.018 (2020).

43 Tajdaran, K., Chan, K., Shoichet, M. S., Gordon, T. & Borschel, G. H. Local delivery of FK506 to injured peripheral nerve enhances axon regeneration after surgical nerve repair in rats. Acta biomaterialia, doi:10.1016/j.actbio.2019.05.058 (2019).

44 Tajdaran, K., Chan, K., Zhang, J., Gordon, T. & Borschel, G. H. Local FK506 dose-dependent study using a novel three-dimensional organotypic assay. Biotechnology and bioengineering 116, 405–414, doi:10.1002/bit.26853 (2019).

45 Soltani, A. M., Allan, B. J., Best, M. J., Mir, H. S. & Panthaki, Z. J. Revision decompression and collagen nerve wrap for recurrent and persistent compression neuropathies of the upper extremity. Annals of plastic surgery 72, 572–578, doi:10.1097/SAP.0b013e3182956475 (2014).

46 Papatheodorou, L. K., Williams, B. G. & Sotereanos, D. G. Preliminary results of recurrent cubital tunnel syndrome treated with neurolysis and porcine extracellular matrix nerve wrap. The Journal of hand surgery 40, 987–992, doi:10.1016/j.jhsa.2015.02.031 (2015).

47 Yeganegi, M., Kandel, R. A. & Santerre, J. P. Characterization of a biodegradable electrospun polyurethane nanofiber scaffold: Mechanical properties and cytotoxicity. Acta biomaterialia 6, 3847–3855, doi:10.1016/j.actbio.2010.05.003 (2010).

48 Wright, M. E. E. et al. Influence of ciprofloxacin-based additives on the hydrolysis of nanofiber polyurethane membranes. Journal of biomedical materials research. Part A 106, 1211–1222, doi:10.1002/jbm.a.36318 (2018).

49 Ma, Z. et al. In vitro and in vivo mechanical properties of human ulnar and median nerves. Journal of biomedical materials research. Part A 101, 2718–2725, doi:10.1002/jbm.a.34573 (2013).

50 Tajdaran, K., Shoichet, M. S., Gordon, T. & Borschel, G. H. A novel polymeric drug delivery system for localized and sustained release of tacrolimus (FK506). Biotechnology and bioengineering 112, 1948–1953, doi:10.1002/bit.25598 (2015).

51 Yalcin, E., Onder, B. & Akyuz, M. Ulnar nerve measurements in healthy individuals to obtain reference values. Rheumatology International 33, 1143–1147, doi:10.1007/s00296-012-2527-9 (2013).

52 Jo, S. et al. Comparing electrical stimulation and tacrolimus (FK506) to enhance treating nerve injuries. Muscle & nerve, doi:10.1002/mus.26659 (2019).

53 Yang, R. K. et al. Dose-dependent effects of FK506 on neuroregeneration in a rat model. Plastic and reconstructive surgery 112, 1832–1840, doi:10.1097/01.prs.0000091167.27303.18 (2003).

54 Bertelli, J. A., Taleb, M., Saadi, A., Mira, J. C. & Pecot-Dechavassine, M. The rat brachial plexus and its terminal branches: an experimental model for the study of peripheral nerve regeneration. Microsurgery 16, 77–85 (1995).

55 Tao, X. et al. Long-term efficacy and side effects of low-dose tacrolimus for the treatment of Myasthenia Gravis. Neurol Sci 38, 325–330, doi:10.1007/s10072-016-2769-5 (2017).

56 Kemper, M. J., Spartà, G., Laube, G. F., Miozzari, M. & Neuhaus, T. J. Neuropsychologic side-effects of tacrolimus in pediatric renal transplantation. Clin Transplant 17, 130–134, doi:10.1034/j.1399-0012.2003.00028.x (2003).

57 Krishna, N., Chiappelli, J., Fischer, B. A. & Knight, S. Tacrolimus-induced paranoid delusions and fugue-like state. Gen Hosp Psychiatry 35, 327.e325-327.e326x, doi:10.1016/j.genhosppsych.2012.07.010 (2013).

58 Santerre, J. P., Woodhouse, K., Laroche, G. & Labow, R. S. Understanding the biodegradation of polyurethanes: from classical implants to tissue engineering materials. Biomaterials 26, 7457–7470, doi:10.1016/j.biomaterials.2005.05.079 (2005).

59 Stroka, K. M., Gu, Z., Sun, S. X. & Konstantopoulos, K. Bioengineering paradigms for cell migration in confined microenvironments. Curr Opin Cell Biol 30, 41–50, doi:10.1016/j.ceb.2014.06.001 (2014).

60 Perez, R. A. & Kim, H. W. Core-shell designed scaffolds for drug delivery and tissue engineering. Acta biomaterialia 21, 2–19, doi:10.1016/j.actbio.2015.03.013 (2015).

61 Li, X. et al. Immunophilin FK506 loaded in chitosan guide promotes peripheral nerve regeneration. Biotechnol Lett 32, 1333–1337, doi:10.1007/s10529-010-0287-8 (2010).

62 Abram, S. E., Yi, J., Fuchs, A. & Hogan, Q. H. Permeability of injured and intact peripheral nerves and dorsal root ganglia. Anesthesiology 105, 146–153, doi:10.1097/00000542-200607000-00024 (2006).

63 Hirakawa, H., Okajima, S., Nagaoka, T., Takamatsu, T. & Oyamada, M. Loss and recovery of the blood-nerve barrier in the rat sciatic nerve after crush injury are associated with expression of intercellular junctional proteins. Experimental cell research 284, 196–210, doi:10.1016/s0014-4827(02)00035-6 (2003).

64 Bouldin, T. W., Earnhardt, T. S. & Goines, N. D. Restoration of blood-nerve barrier in neuropathy is associated with axonal regeneration and remyelination. J Neuropathol Exp Neurol 50, 719–728, doi:10.1097/00005072-199111000-00004 (1991).

65 Liu, Q., Wang, X. & Yi, S. Pathophysiological Changes of Physical Barriers of Peripheral Nerves After Injury. Frontiers in Neuroscience 12, doi:10.3389/fnins.2018.00597 (2018).

66 Dahlin, L. B., Rydevik, B., McLean, W. G. & Sjöstrand, J. Changes in fast axonal transport during experimental nerve compression at low pressures. Experimental neurology 84, 29–36, doi:10.1016/0014-4886(84)90003-7 (1984).

67 Rydevik, B., Lundborg, G. & Bagge, U. Effects of graded compression on intraneural blood blow. An in vivo study on rabbit tibial nerve. The Journal of hand surgery 6, 3–12, doi:10.1016/s0363-5023(81)80003-2 (1981).

68 Percie du Sert, N. et al. Reporting animal research: Explanation and elaboration for the ARRIVE guidelines 2.0. PLOS Biology 18, e3000411, doi:10.1371/journal.pbio.3000411 (2020).

69 Zaimi, A. et al. AxonDeepSeg: automatic axon and myelin segmentation from microscopy data using convolutional neural networks. Scientific reports 8, 3816, doi:10.1038/s41598-018-22181-4 (2018).

70 Richner, M., Jager, S. B., Siupka, P. & Vaegter, C. B. Hydraulic Extrusion of the Spinal Cord and Isolation of Dorsal Root Ganglia in Rodents. Journal of visualized experiments : JoVE, 55226, doi:10.3791/55226 (2017).

71 Daeschler, S. C., Zhang, J., Gordon, T. & Borschel, G. H. Optical Tissue Clearing Enables Rapid, Precise and Comprehensive Assessment of Three-Dimensional Morphology in Experimental Nerve Regeneration Research. bioRxiv, 2021.2001.2028.428623, doi:10.1101/2021.01.28.428623 (2021).

72 Qi, Y. et al. FDISCO: Advanced solvent-based clearing method for imaging whole organs. Sci Adv 5, eaau8355, doi:10.1126/sciadv.aau8355 (2019).

73 Carro, M., Paroutis, P., Woolside, M. & Harrison, R. Improved Imaging of Cleared Samples with ZEISS Lightsheet Z.1: Refractive Index on Demand. (2015). <https://www.google.com/url?sa=t&rct=j&q=&esrc=s&source=web&cd=&cad=rja&uact=8&ved=2ahUKEwiYq8CO6uPtAhU7AGMBHSohDMoQFjAAegQIBhAC&url=https%3A%2F%2Fasset-downloads.zeiss.com%2Fcatalogs%2Fdownload%2Fmic%2Fb91bc689-66ab-4d85-b3f6-c31ebea22ba7%2FEN_wp_Lightsheet-Z.1_Expanding-Clearing-Solutions.pdf&usg=AOvVaw215TogF5NamyFoohy0IlGn>.

74 Daeschler, S. C. et al. Clinically Available Low Intensity Ultrasound Devices do not Promote Axonal Regeneration After Peripheral Nerve Surgery-A Preclinical Investigation of an FDA-Approved Device. Frontiers in neurology 9, 1057, doi:10.3389/fneur.2018.01057 (2018).

